# Human vein-to-artery endothelial cell fate transition is driven by VEGF/ERK activation and PI3K inhibition

**DOI:** 10.64898/2025.12.17.694993

**Authors:** Z. Amir Ugokwe, A.L. Pyke, E. Trimm, M. Chakraborty, X. Fan, L.T. Ang, K.M. Loh, K. Red-Horse

**Affiliations:** Department of Biology, Stanford University, Stanford, CA, USA; Department of Genetics, Stanford University, Stanford, CA, USA; Biophysics Graduate Program, Stanford University, Stanford, CA, USA; Institute for Stem Cell Biology and Regenerative Medicine, Stanford University, Stanford, CA, USA; Department of Urology, Stanford University, Stanford, CA, USA; Department of Developmental Biology, Stanford University, Stanford, CA, USA; Howard Hughes Medical Institute, Stanford University, Stanford, CA, USA

**Keywords:** Angiogenesis, vasculogenesis, vein-to-artery endothelial conversion, cell fate switch, endothelial differentiation, human pluripotent stem cells

## Abstract

Artery endothelial cells (ECs) arise through different pathways, including differentiation from mesodermal cells (vasculogenesis) or from already established vein or capillary plexus ECs (angiogenesis), the latter being most common during embryonic development and regeneration. Understanding the vein-to-artery (v2a) transition could improve revascularization therapies, but progress is limited by a lack of human models. Here, we develop a human pluripotent stem cell (hPSC) differentiation protocol that models the v2a EC conversion. Comparing v2a and mesoderm-to-artery (m2a) transcriptomes with publicly available single cell RNA sequencing (scRNA-seq) data from human embryos showed they reflected angiogenesis- and vasculogenesis-derived artery ECs, respectively. This reductionist system revealed that VEGF activation alongside PI3K inhibition was sufficient for vein ECs to acquire arterial identity within 48 hours. We model a critical step in vascular development and define the minimal signals required for artery differentiation from veins, providing a framework to promote this conversion in revascularization or therapeutic contexts.

## INTRODUCTION

Blood vessels permeate all organs and carry out functions as diverse as regulating oxygen and nutrient exchange, organizing immune surveillance, and facilitating interorgan communication through endocrine signaling (Potente and Mäkinen, 2017; Augustin and Koh, 2024). Given the paramount physiological roles of the vasculature, vascular dysfunction contributes to a broad spectrum of chronic diseases (Pasut *et al*., 2025). Therapeutic interventions aimed at restoring or enhancing vascular function hold promise for improving outcomes, but despite considerable progress in model organisms, critical mechanistic knowledge of human vascular development pathways remain unresolved. Here we introduce an *in vitro* model of the angiogenesis phase of human vascular development wherein vein endothelial cells (vECs) transition to artery ECs (aECs), and we apply this model to explore the signaling pathways regulating the vein-to-artery transition (v2a).

ECs arise in two major waves, denoted vasculogenesis and angiogenesis. First, during vasculogenesis, lateral mesoderm progenitors differentiate into the first ECs; this is unique because it generates the first blood vessels *de novo*. Vasculogenesis occurs in the early embryo (embryonic days ∼7-9 in mice and ∼16-21 in humans; 12-22 hours post fertilization in zebrafish) and generates the first intraembryonic arteries and veins, such as the dorsal aorta and cardinal vein, respectively (Weinstein, 1999; Chong *et al*., 2011; Kohli *et al*., 2013; Lindskog *et al*., 2014; Loh and Ang, 2024). Later in angiogenesis, new blood vessels are generated from pre-existing ones.

An emerging concept in vascular biology is that during angiogenesis, ECs from existing veins sprout and differentiate into capillary and artery ECs during both development and regeneration in adults (Red-Horse *et al*., 2010; Xu *et al*., 2014; Kametani *et al*., 2015; Red-Horse and Siekmann, 2019; Park *et al*., 2021; Trimm and Red-Horse, 2023; Bovay *et al*., 2025). One reason this may be critical is that artery ECs in growing branches typically exhibit low to no proliferation such that expansion of arteries requires recruitment of new ECs from veins, which are generally more proliferative (Fang *et al*., 2017; Jin *et al*., 2017; Sugden *et al*., 2017; Su *et al*., 2018; Luo *et al*., 2021; Bovay *et al*., 2025; Diwan *et al*., 2025). Lineage-tracing experiments in the mouse heart and postnatal retina and live imaging of regenerating zebrafish tissues have demonstrated a directional flow of ECs from veins to capillaries to arteries as the vascular network expands and matures (Chen *et al*., 2014; Xu *et al*., 2014; Pitulescu *et al*., 2017; Su *et al*., 2018; Lee *et al*., 2021; Park *et al*., 2021; Zarkada *et al*., 2021; Bovay *et al*., 2025). Trajectory analysis of single-cell RNA sequencing (scRNA-seq) data suggests that this is a common theme in developing mouse and human embryos (Hou *et al*., 2022; McCracken *et al*., 2022; Phansalkar *et al*., no date). Thus, to fully model human vascular development *in vitro*, we would need a system that recapitulates the vein-to-artery EC lineage conversion.

Recent advances have enabled the generation of ECs from hPSCs (Kubo *et al*., 2005; James *et al*., 2010; Ditadi *et al*., 2015; Zhang *et al*., 2017; Paik *et al*., 2018; McCracken *et al*., 2019; Wang *et al*., 2020; Ang *et al*., 2022; Loh and Ang, 2024; Gong *et al*., 2025; Loh *et al*., 2025), drawing on knowledge from vascular development in animal models. In these differentiation protocols, lateral mesoderm progenitors (i.e., non-ECs) are directly differentiated into ECs (Ditadi *et al*., 2015; Park *et al*., 2018; Rosa *et al*., 2019; Ang *et al*., 2022, 2026; Pan *et al*., 2025), thereby modeling vasculogenesis and *de novo* EC production (Fish and Wythe, 2015; McCracken *et al*., 2023). However, we lack a defined and tractable model of the vein-to-artery EC fate transitions that occur in subsequent stages of vascular development and adult vascular regeneration, namely angiogenesis. This is particularly important, because most organs are vascularized through angiogenesis. Presumably, there could be mechanistic differences between vasculogenesis and angiogenesis, as vasculogenesis entails the *de novo* establishment of EC identity, whereas angiogenesis entails an arteriovenous identity switch in already-established ECs.

To develop an *in vitro* human model system of angiogenesis, we leverage an established protocol for studying human vasculogenesis that entails the rapid and efficient differentiation of hPSCs into lateral mesoderm, and subsequently, >90% pure artery and vein ECs (Ang *et al*., 2022). This homogeneity facilitates mechanistic discovery because genetic and biochemical experiments can be performed with higher precision. Consistent with known *in vivo* signals and timing (Lawson, Vogel and Weinstein, 2002; Potente, Gerhardt and Carmeliet, 2011; Pitulescu and Adams, 2014; Daems *et al*., 2024; Loh and Ang, 2024), this protocol’s mesoderm-to-artery (m2a) EC differentiation step requires activation of VEGF/ERK, TGFβ, and endogenous NOTCH signaling, alongside inhibition of PI3K, BMP and WNT. Mesoderm-to-vein (m2v) EC differentiation entails two steps: in the first step, VEGF/ERK activation generates primed ECs, and in the second step, VEGF/ERK inhibition induces venous identity; both steps require NOTCH inhibition and PI3K activation (Ang *et al*., 2026) **(Fig. 1A)**.

**Figure 1.**
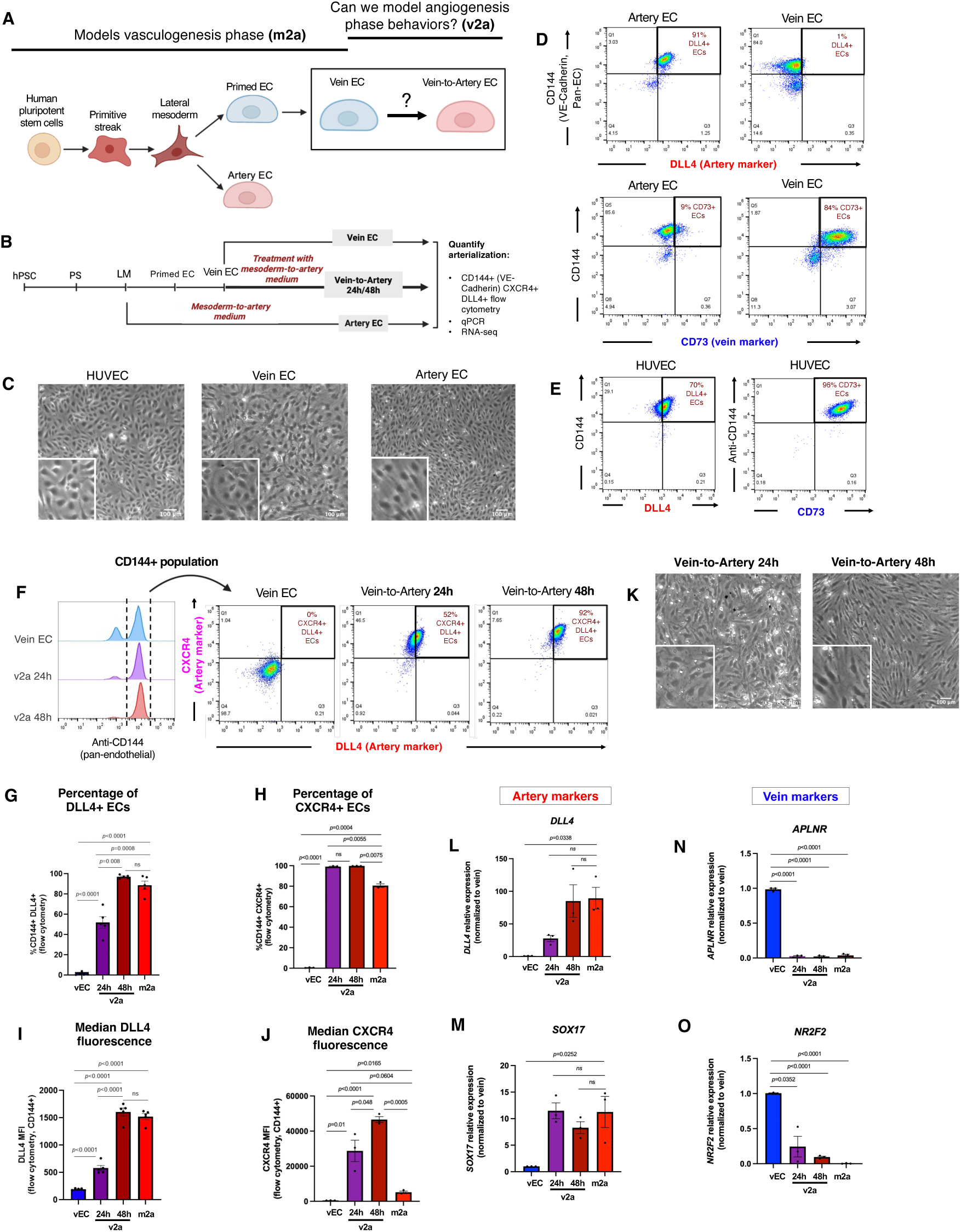
Establishing an in vitro model of the human vein-to-artery EC transition using pluripotent stem cell differentiation. **(A)** Schematic of the differentiation protocol adapted from Ang et al. (Ang *et al*., 2022), showing generation of venous endothelial cells (vECs) from hPSCs. A question mark denotes the experimental extension introduced in this study to model vein-to-artery conversion in vitro. **(B)** Experimental design for assessing arterial ECs conversion in vECs treated with mesoderm-to-artery (m2a) medium for 24 or 48 hours. **(C)** Brightfield images of HUVECs, vECs, and aECs demonstrate preserved endothelial morphology across all states. Scale bars, 100 µm. **(D and E)** Flow cytometry plots showing artery (DLL4), vein (CD73), and pan-EC (CD144; VE-Cadherin) marker expression in hPSC-derived ECs **(D)** and HUVECs **(E)**. **(F)** Representative flow cytometry plots showing arterial markers (DLL4+CXCR4+) within the endothelial (CD144+) cell population in hESC-derived vECs and v2a at 24 and 48 hours. **(G-J)** Percentage of CD144+, DLL4+ ECs **(G)** and CD144+, CXCR4+ ECs **(H)** in the indicated conditions and their corresponding median fluorescence intensities (MFIs)**(I, J),** showing progressive arterial marker induction. **(K)** Brightfield image of v2a ECs at 24 and 48 hours reveals typical endothelial morphology. Scale bar, 100 µm. **(L-O)** qPCR analysis of arterial **(L, M)** and venous **(N, O)** markers in the indicated cells supports transition toward arterial fate and away from venous fate. Data represents 2-5 independent experiments. Statistical significance was determined using unpaired two-tailed t-tests; p-values are shown, “ns” indicates p values > or = 0.05. Error bars are mean +/- SEM. In bar plots, each dot represents one independent experiment.

Here, we build upon the Ang et al. protocol (Ang *et al*., 2022, 2026) to generate vein ECs that models *de novo* vasculogenesis and add a subsequent vein-to-artery (v2a) EC step that models angiogenesis from pre-existing ECs. Vein ECs fully transition to artery ECs after 48 hours in the presence of components that induce the m2a transition. Primed ECs–those emerging from the first step in vein differentiation (Ang *et al*., 2026) –could also differentiate to artery when exposed to the same media, which we denoted as p2a. Comparisons to publicly available scRNA-seq data from human embryos indicated that the m2a transcriptome reflects a vasculogenesis-derived artery path, whereas the v2a and p2a align closer to angiogenesis-derived arterial ECs. This platform to model the human vein-to-artery fate transition during angiogenesis will be useful for studies to discover the molecular mechanisms and genes important for generating arteries from veins, both in development and vascular regeneration.

## RESULTS

### An *in vitro* model of human vein-to-artery endothelial cell lineage conversion

Given that many arteries develop from vein ECs during angiogenesis (Herbert *et al*., 2009; Red-Horse *et al*., 2010; Chen *et al*., 2014; Xu *et al*., 2014; Su *et al*., 2018; Red-Horse and Siekmann, 2019; Lee *et al*., 2021; Hou *et al*., 2022; Phansalkar *et al*., no date), we aimed to generate a human *in vitro* model of vein-to-artery EC fate switching **(Fig. 1A)**. First, we differentiated hPSCs into lateral mesoderm, and subsequently, primed ECs and vein ECs as described previously (Ang *et al*., 2022, 2026), thereby modeling vasculogenesis. We then challenged these vein ECs with the various signals that were previously shown to induce artery EC differentiation directly from mesoderm (Ang *et al*., 2022). Acquisition of arterial identity was assessed by artery marker expression via flow cytometry (e.g., DLL4, CXCR4), qPCR, and bulk population RNA-seq **(Fig. 1B)**.

The Ang et al. protocol (Ang *et al*., 2022) generates highly pure 2D monolayers of either vein or artery ECs from hPSCs, including human embryonic stem cells (hESCs) and induced pluripotent stem cells (hiPSCs), with morphology similar to primary human ECs such as HUVECs **(Fig. 1C)**. Flow cytometry **(Fig. S1A)** demonstrated high purity—most differentiated artery ECs (91%) expressed the artery marker DLL4 with only a few (∼9%) expressing the venous marker CD73 (NT5E) (Ditadi *et al*., 2015; Rosa *et al*., 2019), consistent with arterial identity. Conversely, vein ECs were DLL4-negative (99% of ECs) and CD73-positive (84% of ECs) **(Fig 1D)**. HUVECs anomalously expressed both arterial (DLL4) and venous (CD73) markers, consistent with the widely recognized view that they have ambiguous arteriovenous identity (Aranguren *et al*., 2013) **(Fig. 1E).**

We next tested whether the vein ECs would acquire an arterial fate upon treatment with the six signals that differentiate mesoderm into artery ECs (i.e., m2a medium), which comprises our base medium called “chemically defined medium (CDM2)”(see Methods) with VEGF agonist, TGFβ agonist, PI3K inhibitor, WNT inhibitor, BMP inhibitor, and Vitamin C (Ang *et al*., 2022). To assess arterial conversion, we collected vECs exposed to m2a medium for 24 and 48 hours and quantified the percentage of CD144^+^DLL4^+^CXCR4^+^ cells by flow cytometry, using well-established pan-endothelial (CD144, also known as VE-Cadherin) and arterial-specific (DLL4, CXCR4) markers.

The m2a medium induced ∼50% of the vein EC population to express DLL4 after 24 hours **(Fig. 1F and G)**, whereas ∼100% were CXCR4-positive **(Fig. 1F and H)**. By 48 hours, nearly all (∼100%) ECs–referred to here as v2a–expressed both artery markers, DLL4 and CXCR4 **(Fig. 1F-H).** Quantification of expression levels in ECs showed that both markers increased over 48 hours **(Fig. 1I and J)**. V2a ECs also retained the endothelial morphology observed in other EC states **(Fig. 1K, compared to 1C)**. Similar findings were observed when using hiPSCs for differentiation instead of hESCs **(Fig.** S1B-F).

We next performed qPCR to assess additional vein and artery markers during v2a. *DLL4* expression mirrored protein expression in that it steadily increased until reaching m2a levels at 48 hours **(Fig. 1L)**. However, by 24 hours, the artery-enriched SoxF factor, *SOX17* (Clarke *et al*., 2013; Corada *et al*., 2013), was expressed at m2a levels **(Fig. 1M)**. The kinetics of *SOX17* and *DLL4* expression (i.e. *SOX17* upregulation prior to that of *DLL4*) are consistent with findings in zebrafish and mice that SoxFs bind to, and activate, *Dll4* enhancers during vascular development (Sacilotto *et al*., 2013; McCracken *et al*., 2023). Both vein-enriched genes queried, *NR2F2* and *APLNR*, were almost completely turned off after 24 hours **(Fig 1N and O)**.

In sum, these findings demonstrate that the factors inducing human artery EC differentiation directly from mesoderm *in vitro* can also arterialize already established vein ECs. This suggests that the common *in vivo* vein-to-artery pathway occurring during angiogenesis can be modeled *in vitro*.

### Vein-to-artery endothelial cells lose vein identity and exhibit a global transcriptional transition toward artery fate

To characterize the vein-to-artery transition at the global transcriptional level, we performed bulk RNA sequencing (RNA-seq) on CD34*^+^* v2a ECs at 24 and 48 hours, as well as vECs and aECs generated directly from lateral mesoderm **(Fig. 1B)**. Quality control analyses using the global transcriptomes to calculate Euclidean distance and perform principal component analysis (PCA) revealed tight clustering of biological replicates **(Fig. S2A and B).** Upon 24 and 48 hours of exposure to arterializing media, 1345 and 2590 genes, respectively, became differentially expressed relative to the starting vein ECs (adjusted p-value < 0.05, |log2 fold-change| > 2; **Fig. 2A**, **Table S1)**. At 48 hours of vein-to-artery conversion, arterial genes *CXCR4, NOTCH1/4,* and *HEY1* were upregulated, whereas venous markers *NR2F2, APLNR, EMCN,* and *STAB2* were downregulated **(Fig. 2B)**.

**Figure 2.**
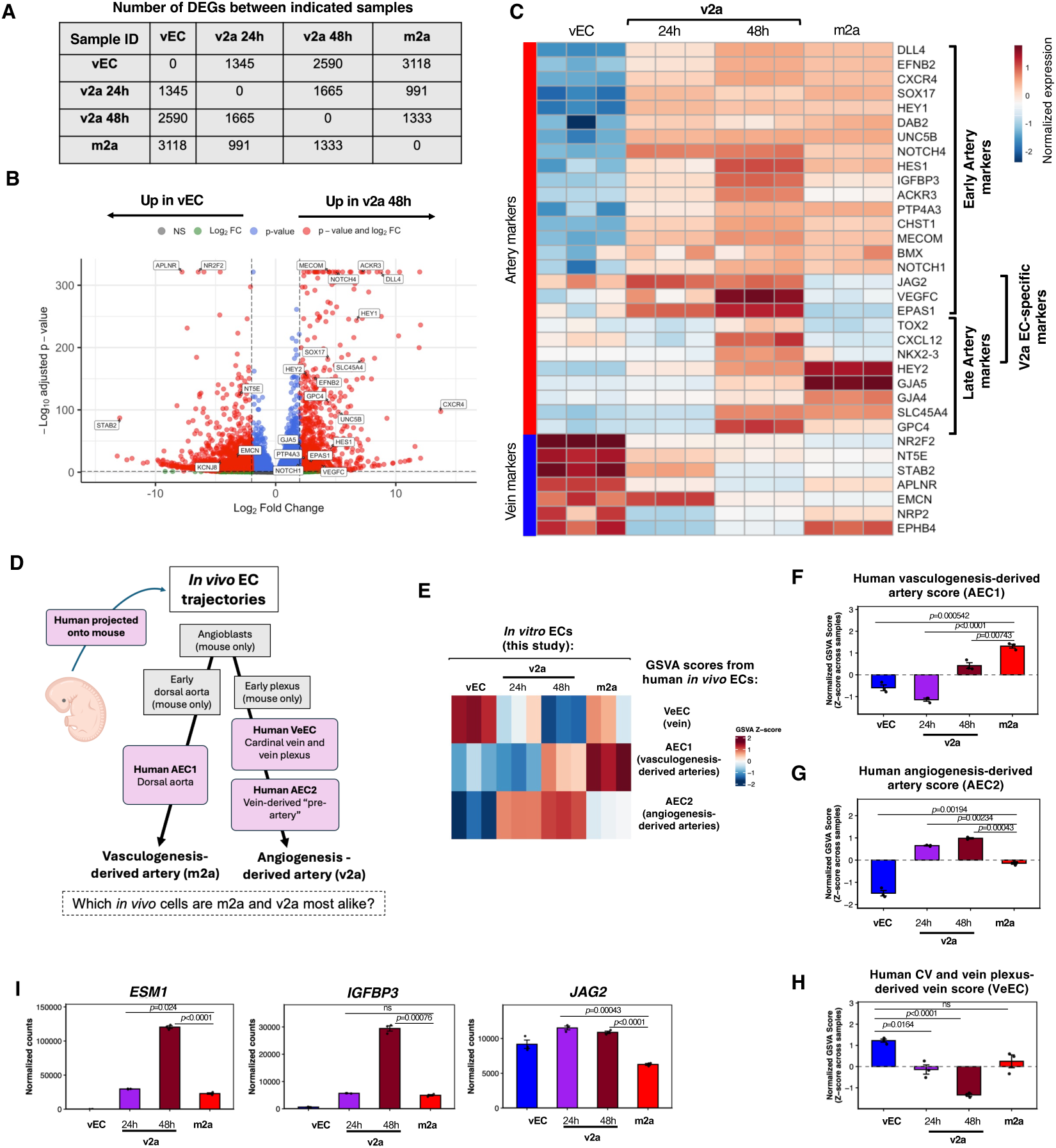
The vein-to-artery protocol generates ECs with a transcriptional profile aligning with their *in vivo* correlates—human vein-derived arteries. **(A)** Pairwise comparisons of number of differentially expressed genes (DEGs) between the indicated cells. (DEGs were defined as those with adjusted p < 0.05 and |log2 fold change|>2) **(B)** Volcano plot comparing gene expression differences between v2a 48h and vECs with labeling of select canonical arterial and venous genes. **(C)** Heatmap of arterial and venous marker genes, including novel genes described in Su et al. 2018, showing a gradual loss of venous identity and arterial induction with the v2a protocol. **(D)** Schematic of *in vivo* cell trajectories along the vasculogenic and angiogenic artery development pathways in early mouse and human embryos as shown by scRNA-seq in Hou et al., 2021. Individual cell clusters along the trajectories are shown in grey and pink boxes. **(E-H)** Heatmaps **(E)** and bar plots **(F-H)** showing how bulk RNA sequencing-derived transcriptional profiles from indicated in vitro ECs compared to GSVA gene scores from human vasculogenic-derived arterial (AEC1), angiogenic-derived arterial (AEC2) and venous (VeEC), which were calculated from cluster-defining genes in scRNA-seq clusters shown in D. **(I)** DESeq2 normalized expression levels of select genes that define *in vivo* angiogenesis-derived artery ECs and are expressed in v2a over m2a. Arterial ECs derived from the lateral mesoderm (see schematic in Fig. 1B) are labeled “m2a” in panels **(B–I)**. In bar plots, each dot represents one independent cell culture well, and error bars are mean +/- SEM. Pairwise comparisons were performed using Welch’s two-sample t-tests. p-values are shown, “ns” indicates p values > and/or = 0.05.

Assessing well established vein and artery markers showed a few notable patterns (Su *et al*., 2018) **(Fig. 2C)**. First, most markers were changed in a manner consistent with v2a ECs being fully arterial after 48 hours. Second, artery markers were categorized into either early (up at 24 hours) or late (up at 48 hours) induction, as seen previously in coronary vein-to-artery development in the mouse (Su *et al*., 2018). Third, some artery markers were only upregulated in v2a ECs and not m2a ECs, some of which were novel artery genes that we discovered were upregulated during vein-to-artery angiogenesis in the developing heart (e.g., *TOX2, CXCL12, JAG2, VEGFC, EPAS1*) (Su *et al*., 2018).

We next tested whether our *in vitro* differentiation recapitulated *in vivo* artery development via vasculogenesis and/or angiogenesis. Hou et al. (Hou *et al*., 2022) generated, and validated with lineage tracing, a scRNA-seq dataset of ECs progressing along both the vasculogenesis and angiogenesis trajectories in the mouse, which primarily reflects the early vasculature containing the dorsal aorta, cardinal vein, and early vascular plexuses. They then used the mouse trajectory to integrate human embryonic scRNA-seq data from analogous stages (Hou *et al*., 2022) **(Fig. 2D)**. To investigate how our *in vitro* EC populations compared to the human-derived vasculogenesis and angiogenesis arterial trajectories, we used Gene Set Variation Analysis (GSVA) (Hänzelmann, Castelo and Guinney, 2013) to generate enrichment scores for each of the human *in vivo* cell clusters shown in **Fig. 2D**. These GSVA scores were then calculated for our *in vitro* differentiated cells. Strikingly, *in vitro* m2a ECs have a higher *in vivo* human vasculogenesis-derived artery score (m2a; mean z-score=1.4), relative to the angiogenesis-derived artery scores (v2a 24h and 48h; mean z-score= -0.8, and 0.4, respectively) **(Fig. 2E and F)**. Conversely, *in vitro* v2a ECs have a higher human angiogenesis-derived artery score (v2a 24h and 48h; mean z-score= 0.6 and 1.0, respectively) than vasculogenesis-derived artery score (m2a; mean z-score=-0.1) **(Fig 2E and G).** Only *in vitro* vein ECs were highly scored with the *in vivo* vein-derived signature **(Fig 2E and H)**. These data support the notion that the m2a step of the *in vitro* differentiation reflects vasculogenesis, while the v2a step models angiogenesis.

While inspecting the genes driving the *in vivo* GSVA scores, there were a few notable markers **(Fig. 2I and S3)**. For instance, v2a ECs showed elevated levels of the angiogenesis-derived artery gene *endothelial cell-specific molecule 1* (*Esm1)*. Esm1 binds to fibronectin, competing with VEGF for fibronectin binding, consequently displacing VEGF and increasing VEGF bioavailability and signaling through VEGFRs (Rocha *et al*., 2014). It has also been established as a marker of angiogenesis-derived pre-artery cells *in vivo*: in the postnatal retina, *Esm1* is a specific marker of pre-artery tip cells and lineage tracing shows they migrate into mature arteries (Xu *et al*., 2014; Pitulescu *et al*., 2017; Stewen *et al*., 2024). *Esm1* also identifies angiogenesis-derived pre-artery ECs in both the heart and intestine (Su *et al*., 2018; Cano *et al*., 2024; Bovay *et al*., 2025). Two other genes, *Igfbp3* and *Jag2* that significantly contribute to angiogenesis-derived artery scores *in vivo* were found to be highly expressed in v2a ECs, and they have previously been identified as markers of vein-derived artery ECs in the heart (Su *et al*., 2018). Together, these data suggest that vasculogenesis- and angiogenesis-derived arteries have both shared and distinct genetic regulators.

### Defining the extracellular signals that drive the human vein-to-artery EC transition

Inspired by the observation that v2a ECs express some unique artery genes when compared to m2a ECs, we considered whether they required different signaling pathways to attain arterial fate. Indeed, vasculogenesis (*de novo* EC specification) and angiogenesis (which entails an arteriovenous identity shift in pre-existing ECs) could be mechanistically different. To this end, we individually withheld each component of m2a arterializing medium (i.e., VEGF agonist, TGFβ agonist, PI3K inhibitor, WNT inhibitor, or BMP inhibitor were individually withheld) during vein-to-artery differentiation for 24 hours **(Fig. 3A)**.

**Figure 3.**
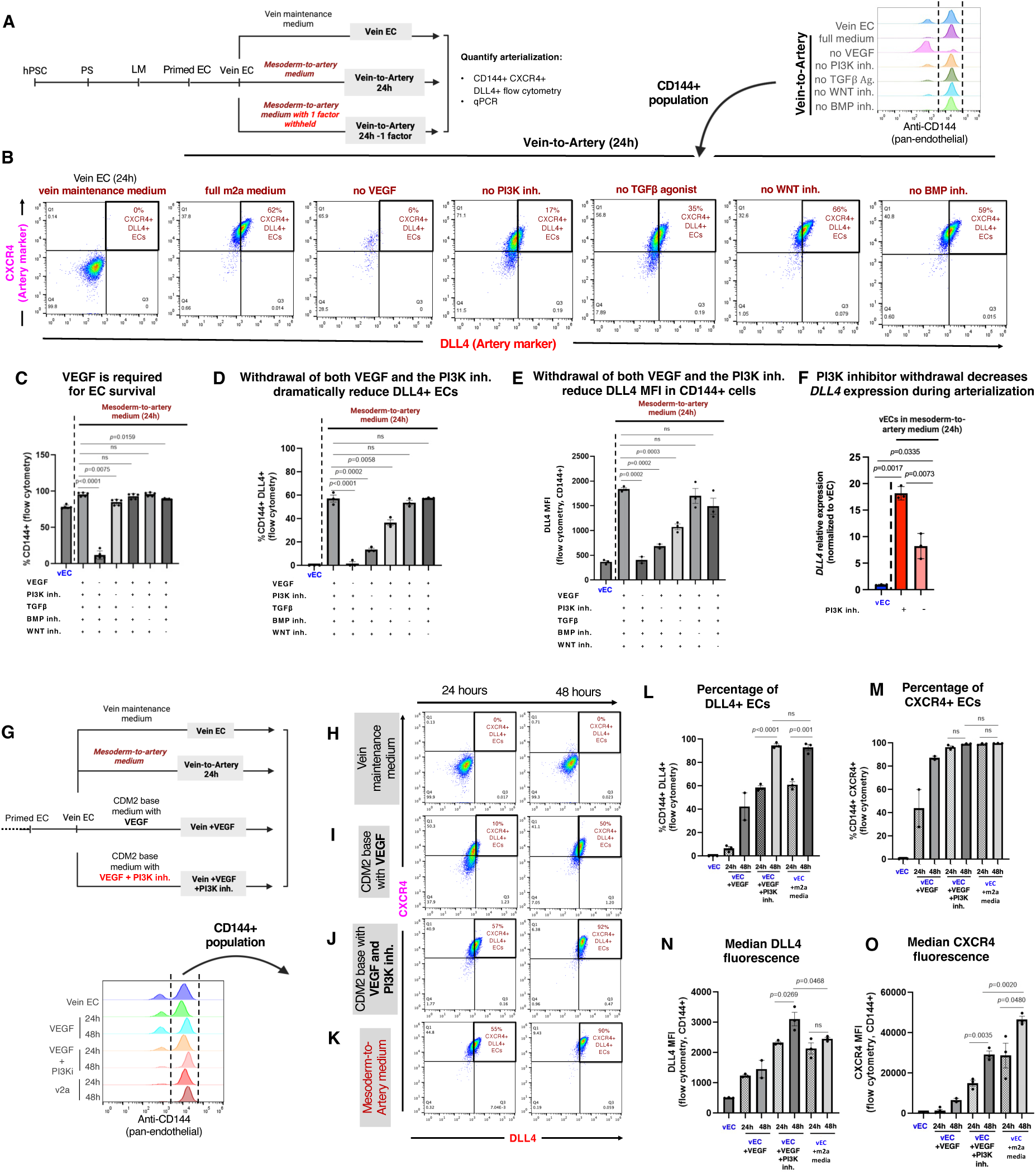
PI3K inhibition and VEGF are sufficient to arterialize vein endothelial cells. **(A)** Strategy to identify individual factors in m2a medium required for vein arterialization **(B)** Representative flow cytometry plots of CD144-gated cells showing percentage of DLL4^+^CXCR4^+^ ECs when the indicated factor is withheld from m2a medium for 24 hours. **(C)** Quantification of CD144^+^ cells in live populations, showing VEGF removal reduces EC numbers. **(D, E)** Quantification of DLL4 within CD144^+^ cells: **(D)** percentage of CD144^+^DLL4^+^ cells and **(E)** DLL4 MFI across conditions. **(F)** qPCR of *DLL4* within CD144^+^ v2a ECs cultured 24 h in PI3K inhibitor-withheld media, normalized to CD144 and relative to vECs. **(G)** Experimental design schematic: vECs cultured 24 h in (1) vein maintenance medium, (2) m2a medium, (3) CDM2 base medium + VEGF, or (4) CDM2 base medium + VEGF + PI3K inhibitor. **(H-J)** Representative flow cytometry plots of CD144^+^ DLL4^+^CXCR4^+^ cells across four treatment conditions. **(K-N)** Quantification of CD144^+^ DLL4^+^ **(K)** and CD144^+^CXCR4^+^ **(L)** percentages, with corresponding **(M)** DLL4 and **(N)** CXCR4 MFIs. Each dot represents a biological replicate. In bar plots, individual dots represent independent biological replicates. Error bars are mean +/- SEM. Statistical significance was determined by unpaired two-tailed *t*-test; *p*-values are shown, “ns” indicates p values > or = 0.05.

Flow cytometry analysis revealed that removal of VEGF resulted in a low proportion of ECs in the culture, **(Fig. 3B and C)**. Only ∼10% of cells expressed CD144 **(Fig. 3C)** and almost none of these expressed DLL4 **(Fig. 3D and E)**, consistent with longstanding observations that VEGF is required for EC survival in other contexts (Carmeliet *et al*., 1996; Lawson, Vogel and Weinstein, 2002; Coultas, Chawengsaksophak and Rossant, 2005; Casie Chetty *et al*., 2017). Removal of the PI3K inhibitor (GDC-0941) also had a dramatic, but distinct effect. Most of the cells remained endothelial **(Fig. 3C)**, but those that were DLL4^+^ dropped from ∼60% to ∼15% **(Fig. 3D)** and the few DLL4+ cells expressed DLL4 at half the levels of cells in full m2a medium conditions **(Fig. 3E)**. Removing the TGFβ agonist had an effect similar to PI3K inhibitor, but the magnitude of the effect was much milder **(Fig. 3B-E)**. In contrast, removal of either WNT inhibitor (XAV939) or BMP inhibition (DMH1) did not significantly affect the number of DLL4+ ECs **(Fig. 3C and D)** or its expression level **(Fig. 3E)** at 24 hours. Taken together, VEGF/ERK activation and PI3K inhibition appear to be the major drivers of the vein-to-artery EC conversion. While VEGF activation and PI3K inhibition are required for the induction of arterial markers (e.g., DLL4), VEGF is additionally required for EC survival and/or maintenance of pan-EC identity.

To further validate the importance of PI3K inhibition on arterial identity, we performed qPCR comparing v2a ECs cultured with or without PI3K inhibitor. Aligning with protein levels, *DLL4* transcripts were significantly reduced by approximately half in the absence of PI3K inhibition **(Fig. 3F)**.

Together, these data demonstrate that while BMP inhibitor and WNT inhibitor promote the generation of artery ECs from mesoderm (Ang *et al*., 2022), they are not required for the transition of vein ECs into artery ECs. Inhibition of BMP and WNT is crucial to prevent lateral mesoderm cells from transitioning into cardiac and other types of mesoderm (Ang *et al*., 2022). However, during angiogenesis, cells have already acquired EC identity and presumably there is no need to inhibit the alternative lineages that mesoderm cells can access, such as cardiac precursors

### VEGF activation and PI3K inhibition are sufficient to promote arterialization of vein endothelial cells

To test whether VEGF activation and PI3K inhibition are sufficient to induce arterial identity, vein ECs were cultured for 24 and 48 hours under four conditions: (1) vein maintenance medium (consisting of EGM2 + SB505124 (2 µM) + RO4929097 (2 µM), as previously described (Ang *et al*., 2022)), (2) m2a medium (our base medium called CDM2 plus artery factors described in above section), (3) CDM2 base medium (Ang *et al*., 2022) supplemented with VEGF, and (4) CDM2 base medium supplemented with VEGF together with the PI3K inhibitor (GDC-0941)**(Fig. 3G)**. As a negative control, there was minimal DLL4 and CXCR4 expression in CD144^+^ cells maintained in vein maintenance medium **(Fig. 3H)**. Addition of VEGF increased the percentage of CD144^+^DLL4^+^CXCR4^+^ cells to 10% and 50% **(Fig. 3I, L, and M)** at 24 and 48 hours, respectively, and increased expression of DLL4 and CXCR4 protein **(Fig. 3N and O)**, but at levels far below those induced by m2a medium (55%–90% DLL4^+^CXCR4^+^ ECs)**(Fig 3K)**. In contrast, co-treatment with VEGF and GDC-0941 steadily increased the percentage of the CD144^+^DLL4^+^CXCR4^+^ population over 48 hours to 92%, reaching levels similar to those in full m2a medium (90%; **Fig. 3J-O)**. Despite this similarity in population frequency, DLL4 protein levels were significantly higher in VEGF + GDC-0941-treated vein ECs, showing a 1.3-fold increase compared to v2a ECs at 48 hours **(Fig. 3N).** In contrast, CXCR4 expression was reduced in VEGF + GDC-0941-treated vein ECs relative to v2a ECs at the same timepoint, showing a 1.7-fold decrease **(Fig. 3O)**. Similar results were obtained with iPSCs **(Fig. S4)**. These results are consistent with previous reports that PI3K inhibition promotes artery identity in ECs (Hong *et al*., 2006; Ditadi *et al*., 2015; Ang *et al*., 2022; Sabata *et al*., 2025).

These data show that once vein ECs are made from mesodermal cells, they can be transitioned to artery identity with minimal initiating signals, namely simultaneous VEGF addition and GDC-0941.

### Primed endothelial cells also arterialize through an angiogenic-like pathway

Given that *in vivo* angiogenic arterialization often proceeds through intermediate primed or capillary-like states (Red-Horse *et al*., 2010; Xu *et al*., 2014; Red-Horse and Siekmann, 2019; Park *et al*., 2021; Trimm and Red-Horse, 2023; Bovay *et al*., 2025; Pan *et al*., 2025), we next asked whether earlier primed ECs are similarly competent to arterialize and how this route compares to vasculogenesis- versus angiogenesis-derived arterial trajectories. To this end, hPSC-derived primed ECs were cultured in m2a medium for 24 or 48 hours; we denoted this primed EC-to-artery conversion system “p2a” **(Fig. 4A)**. Notably, primed ECs exposed to arterializing cues more rapidly upregulated DLL4 induction when compared to v2a. In the p2a system, nearly all CD144^+^ ECs became DLL4^+^ at 24 hours **(Fig. 4B and C),** with expression levels even surpassing m2a and v2a ECs **(Fig. 4D)**.

**Figure 4.**
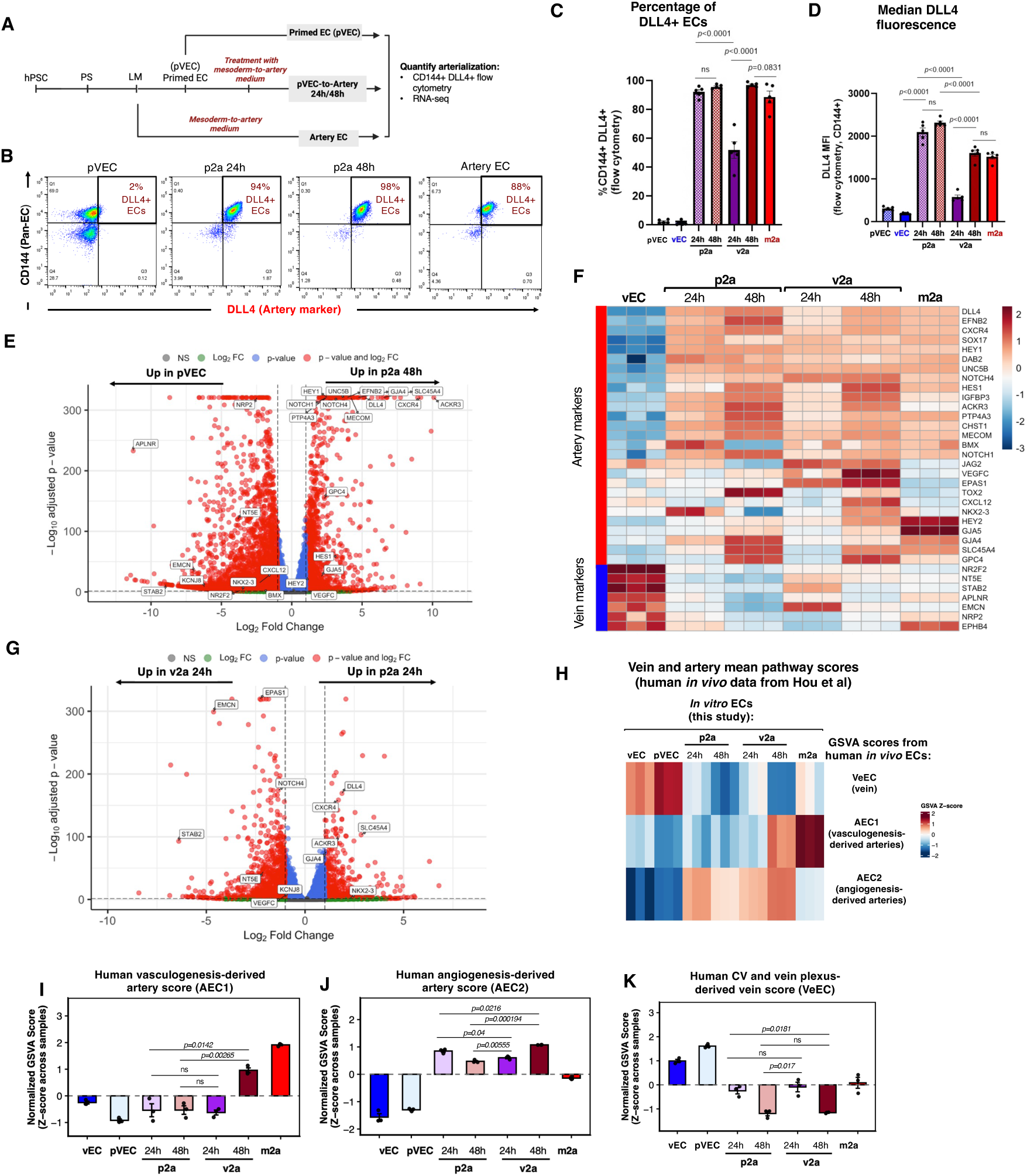
Primed ECs are capable of arterializing and mimic angiogenesis-derived arteries *in vivo*. **(A)** Schematic of primed EC-to-artery (p2a) protocol **(B)** Flow cytometry of CD144+ ECs showing more rapid induction of DLL4 in primed ECs (pVECs) compared with vECs. **(C)** Quantification of the percentage of DLL4+CD144+ cells at 24h and 48h. **(D)** Percentage of DLL4+ ECs and MFI of DLL4, indicating partial acquisition of arterial marker expression in p2a cells. **(E)** Volcano plot of differential expression analysis of arterial and venous genes in p2a 48h vs untreated pVECs. **(F)** Heatmap of artery and vein marker genes across our in vitro conditions, demonstrating more robust arterial gene induction in p2a 24h compared with v2a 24h. **(G)** Volcano plot showing a direct comparison of p2a 24h and v2a 24h highlighting downregulation of venous genes and upregulation of arterial-associated genes in p2a cells. **(H-K)** Heatmap **(H)** and bar plots **(I-K)** showing how bulk RNA sequencing-derived transcriptional profiles from indicated *in vitro* ECs compare to GSVA gene scores from human vasculogenic-derived arterial (AEC1), angiogenic-derived arterial (AEC2) and venous (VeEC), which were calculated from cluster-defining genes in scRNA-seq clusters shown in Fig. 2D. In bar plots, each dot represents one independent cell culture well, and error bars are mean +/- SEM. Statistical significance for selected pairwise comparisons was determined using unpaired two-tailed *t-tests*. *p*-values are shown, “ns” indicates p values > or = 0.05.

Bulk population RNA-sequencing further validated arterial conversion at the transcriptomic level **(Table S1)**. Differential expression analysis showed strong upregulation of arterial genes (e.g. *HEY1, DLL4, CXCR4*) and downregulation of venous markers (e.g. *NR2F2, NT5E, NRP2*) in p2a ECs 48h relative to untreated primed ECs **(Fig. 4E)**. A heatmap of artery and vein marker genes across all conditions demonstrated that multiple canonical artery markers were strongly induced with the p2a path in patterns similar to v2a, although there were differences for a few genes **(Fig. 4F)**.

We next performed the same *in vivo* comparisons with the angiogenic- and vasculogenic-derived arterial gene sets used to compare v2a cells in **Fig. 2D-H**. p2a vasculogenesis scores were even lower than v2a ECs (-0.5 vs -0.2 at 48h, **Fig. 4H and I)** while angiogenesis scores were approaching those of v2a (0.6 vs 0.8 at 48h, **Fig. 4H and J)**. Like v2a, p2a vein scores were low (-0.5 vs -0.7 at 48h, **Fig. 4H and K)**. These data indicate that while primed ECs can differentiate into vein ECs upon VEGF/ERK inhibition (Ang *et al*., 2026), primed ECs can also form artery ECs in the presence of arterializing signals via an angiogenic mechanism.

## DISCUSSION

Here we present an *in vitro* model to study the venous-to-artery fate conversion in human ECs. This is significant because most methods that generate ECs from hPSCs follow a vasculogenic pathway, whereby ECs are directly generated *de novo* from mesoderm (Ditadi *et al*., 2015; Park *et al*., 2018; Rosa *et al*., 2019; Ang *et al*., 2022, 2026; Pan *et al*., 2025). We introduce an *in vitro* model of angiogenesis whereby vein ECs transition to artery ECs, which is the predominant mode of vascularization during organ development and regeneration (Red-Horse *et al*., 2010; Xu *et al*., 2014; Red-Horse and Siekmann, 2019; Park *et al*., 2021; Trimm and Red-Horse, 2023; Bovay *et al*., 2025). For example, vein ECs exposed to arterial-inductive cues transcriptionally lost venous identity and acquired arterial identity within 48 hours. Canonical arterial markers such as *DLL4* and *CXCR4* were upregulated, while venous genes like *NR2F2* and *APLNR* were repressed. Inhibiting PI3K and activating VEGF signaling was sufficient to override venous identity and drive arterial conversion.

This reductionist *in vitro* system will be an important complement to *in vivo* work, allowing us to draw conclusions about the signaling pathways driving the vein-to-artery transition that can be difficult to achieve *in vivo*. First, *in vivo* mapping and loss- and gain-of-function analyses can identify which cellular and molecular pathways discovered in other organisms influence human EC development. This is important if these pathways are to be translated into clinical treatments. Second, the system can precisely dissect necessity and sufficiency. Our data revealed that once cells are ECs, fewer cues are required to specify arterial fate than during mesoderm-to-EC differentiation. Arterialization of vECs required a minimal regimen of VEGF for survival and PI3K inhibition for artery fate induction. Cell state-dependent responses were also observed. The same signaling factors induced distinct arterial gene programs when applied to pre-differentiated vECs compared with its effects on mesoderm (e.g., v2a-specific markers shown in Fig. 2C). This *in vitro* model will be an invaluable tool for discovering and dissecting the genes driving human artery identity and may help guide future approaches to developing therapies for regenerating arteries.

A key question in the field of stem cell biology is how faithfully *in vitro* differentiation protocols mimic *in vivo* events. The first intraembryonic blood vessels in developing embryos—the dorsal aorta and cardinal vein—form by coalescence of artery and vein ECs that emerge *de novo* from lateral mesoderm. Following this *de novo* formation, embryos generally use sprouting angiogenesis from pre-formed vessels, frequently veins, to generate new organ beds (Red-Horse *et al*., 2010; Xu *et al*., 2014; Red-Horse and Siekmann, 2019; Park *et al*., 2021; Trimm and Red-Horse, 2023; Bovay *et al*., 2025). To investigate which path our m2a and v2a ECs were most similar to, we leveraged a study that generated lineage tracing-validated scRNA-seq datasets capturing both the vasculogenic and angiogenic routes to the arterial lineage in mouse embryos (Hou *et al*., 2022). In this study, the authors then integrated scRNA-seq data from early human embryos providing human gene signatures for both trajectories. Using the bulk sequencing data from our *in vitro* m2a and v2a ECs, we measured how closely these ECs mimicked *in vivo* vasculogenesis and angiogenesis at the transcriptional level. The m2a ECs—reflecting the products of current hPSC differentiation protocols (Ditadi *et al*., 2015; Park *et al*., 2018; Rosa *et al*., 2019; Ang *et al*., 2022, 2026; Pan *et al*., 2025)—were highly similar to vasculogenesis-derived human artery ECs with very low scoring of the angiogenesis and vein signatures. By contrast, the v2a ECs generated in this study were most similar to angiogenesis-derived artery ECs but also exhibited a positive score for vasculogenesis-derived arteries. These data align with our analysis of known canonical artery markers where a prominent difference between m2a and v2a ECs was the additional expression of a set of artery markers in v2a on top of their shared canonical markers.

When assessing the genes driving these patterns, interesting candidates emerged. For example, v2a ECs expressed higher levels of *ESM1*, which is a well-established marker of angiogenesis-derived artery cells *in vivo*. *Esm1* is expressed in pre-artery tips cells in the postnatal retina, and *Esm1CreER*-mediated lineage labeling of these cells traces into mature arteries (Xu *et al*., 2014; Pitulescu *et al*., 2017; Stewen *et al*., 2024). It similarly marks angiogenesis-derived pre-artery ECs in the heart and intestine (Su *et al*., 2018; Cano *et al*., 2024; Bovay *et al*., 2025). *JAG2* and *IGFBP3* are two genes driving *in vivo* angiogenesis-derived artery scores in this study that are highly expressed in v2a ECs and previously identified as marking vein-derived artery ECs in the heart(Su *et al*., 2018). It will be interesting in future studies to interrogate the requirement of all the score-driving genes reported here during human artery development, perhaps through CRISPR screening. In total, the *in vivo* comparisons performed in this study provided evidence that our *in vitro* EC differentiations recapitulate early differentiation events in human embryos.

Our experiments revealed that VEGF activation and PI3K inhibition are sufficient for vein EC arterialization, while inhibition of BMP and WNT signaling, which are required for m2a differentiation, was dispensable. This is likely because WNT and BMP inhibition during mesodermal differentiation prevents alternative fates, such as cardiac lineages, in mesoderm cells (Ang *et al*., 2022), whereas established ECs no longer require suppression of those alternate mesoderm lineage outcomes. While activating VEGF and inhibiting PI3K were crucial to install arterial identity during v2a differentiation, they each had different roles. VEGF signaling was required for EC survival since the number of CD144 cells plummeted upon omitting VEGF from m2a media. VEGF was also required for arterial fate induction as we noticed an increase in expression of DLL4 and CXCR4 with the addition of VEGF alone in the m2a media. In contrast, PI3K inhibition was instead required for ECs to take on arterial fate since its omission from m2a medium resulted in similar numbers of ECs but with vastly decreased levels of DLL4 and CXCR4. This model system will be useful in future studies to define the specific role of different signals since the complexity of *in vivo* studies can make it difficult to separate primary and secondary effects during mechanistic interrogations.

Opposing roles for VEGF/ERK and PI3K signaling for artery and vein EC have been observed *in vivo*. ERK is activated (i.e., phosphorylated) in the dorsal aorta, but not the cardinal vein, in both mouse and zebrafish (Corson *et al*., 2003; Hong *et al*., 2006). During zebrafish vasculogenesis, ERK inhibition blocked artery EC specification. In contrast, inhibiting PI3K activity suppressed the *gridlock/Hey2* mutatation, which impairs arterial differentiation in the developing zebrafish (Hong *et al*., 2006) .The effects of PI3K signaling in the vasculature are mediated by AKT (Hong *et al*., 2006; Kobialka *et al*., 2022), and a constitutively active form of AKT drives ECs towards a venous fate during vasculogenesis (Hong *et al*., 2006). During angiogenesis-mediated artery development in the retina, expression of a constitutively-active form of PI3K in *Esm1*-positive tip cells blocked their ability to differentiate into artery ECs (Sabata *et al*., 2025). This block could be overcome by deleting *NR2F2*, a transcription factor central to vein development. These observations, together with our own, support a model in which PI3K suppression is a key driver of arterial fate acquisition in multiple contexts.

In summary, these findings may have implications beyond development. Since its discovery as the major extracellular signal that controls a myriad blood vessel properties (Fong *et al*., 1995; Shalaby *et al*., 1995; Carmeliet *et al*., 1996; Ferrara *et al*., 1996), VEGF-based therapies have been in development to treat ischemic tissues. However, clinical success has been limited with trials rarely showing long-lasting benefits, possible due to insufficient delivery and lack of generating stable arteries (Ylä-Herttuala *et al*., 2017; Khachigian *et al*., 2023). Our data suggest that stimulating vein ECs with VEGF may not be enough. Instead, arterial specification likely requires both angiogenic signals and suppression of pathways like PI3K that promote venous fate. This dual strategy could improve design of therapies aimed at rebuilding arteries in vascular disease. Additionally, the vein-to-artery EC transition may be more difficult to achieve in adult tissues. For example, coronary artery bypass grafts derived from the saphenous vein, the source in >95% of procedures, fail more readily (>50% fail by 10 years)(Gaudino *et al*., 2017; Royse *et al*., 2022). It has been proposed that this failure is due to the vein’s inability to undergo arterialization (Kudo *et al*., 2007; Berard *et al*., 2013; Lee *et al*., 2021). Future studies could add the factors discovered here to mouse vein-to-artery grafting models and test for improvement.

### Limitations of the study

It is important to clarify that in our study we focused on the cell fate conversion component of angiogenesis, as opposed to sprouting. However, unlike *in vivo* models, our *in vitro* system allowed us to decouple these processes, therefore enabling us to study the gene changes that occur in the vein-to-artery endothelial switch. While our hPSC-based differentiation system models the vein-to-artery EC lineage conversion—a key facet of angiogenesis—*in vitro*, a few limitations should be considered. First, while the transcriptional profiles of our m2a and v2a ECs mirror vasculogenesis- and angiogenesis-derived arterial ECs *in vivo*, respectively, our *in vitro* model does not fully recapitulate the complex *in vivo* process. Second, our study shows that VEGF activation and PI3K inhibition are sufficient to endow vein ECs with arterial identity, but does not incorporate shear stress, which is a known regulator of arterial fate (Buschmann *et al*., 2010; Fang *et al*., 2017; Hwa *et al*., 2017). However, a modular *in vitro* system like ours is amenable to application of additional cues, such as shear stress imposed by microfluidic devices (Sivarapatna *et al*., 2015; Arora *et al*., 2019; Huang *et al*., 2019), to examine their additional contributions. Third, in this study we employ bulk RNA-seq to transcriptionally profile v2a, p2a and m2a EC populations, but further work could incorporate proteomics or scRNA-seq to determine potential cell-to-cell differences. Fourth, an important future step will be to test the molecular mechanisms through which VEGF activation and PI3K inhibition promote arterial identity. We have also identified several genes (i.e., *ESM1*, *JAG2*, and *IGFBP3*) that are enriched in v2a differentiation, relative to m2a differentiation, and genetic perturbations could shed light on their possible functions. Finally, our ECs map onto the early vasculature of before organ-specificity arises (Lin *et al*., 2026). It will be useful in the future to attain tissue specific specialization *<u>in vitro</u>*. Nevertheless, our *in vitro* differentiation system to convert human vein ECs into artery ECs at high purity and scale provides an ideal experimental platform for these, and other, mechanistic investigations.

## ACKNOWLEDGMENTS

This work was previously supported by the NIH T32GM007276 (Z.A.U.; CMB Training Program at Stanford), the Stanford Chem-H Chemistry/Biology Interface (Z.A.U.; CBI) Program, and the NIH R01HL128503 (K.R.-H.). Current funding is provided by the American Heart Association (AHA) Predoctoral Fellowship (Z.A.U.; #25PRE1375905). K.R.-H. is also a Howard Hughes Medical Institute Investigator.

Additional support was provided by Breakthrough T1D Northern California Center of Excellence (K.M.L.), Additional Ventures Expansion Award (K.M.L., L.T.A.), Siebel Stem Cell Institute (L.T.A.), Ludwig Cancer Research (K.M.L.), Stanford Maternal and Child Health Research Institute Chambers Family Foundation Innovation Grant (K.M.L., L.T.A., K.R.H.), Amaranth Foundation (K.M.L.), and Anonymous and Gilbert families (K.M.L.).

L.T.A. is an Additional Ventures Catalyst to Independence Fellow and a Bladder Cancer Advocacy Network Career Development Awardee. K.M.L. is a Human Frontier Science Program Young Investigator (RGY0069/2019), Packard Foundation Fellow, Pew Scholar, Baxter Foundation Faculty Scholar, and The Anthony DiGenova Endowed Faculty Scholar.

We would like to thank Sherry Li Zheng (Kyle Loh’s lab) for providing reagents, and guidance on technical questions. We thank Sawan Jha (Red-Horse lab) for his input and general advice throughout the project. We thank Jeff Naftaly (Red-Horse lab) for providing guidance on quantitative PCR methodology. We thank Qingqing Yin (Kyle Loh’s lab) for kindly sharing details of the artery flow cytometry panel that informed our experiments. We acknowledge Juan Carlos Alcocer (Red-Horse lab), our lab manager, for assistance with ordering and laboratory logistics.

## AUTHOR CONTRIBUTIONS

Conceptualization and methodology, Z.A.U., K.R.-H.,K.M.L., and L.T.A.; investigation, Z.A.U.; data analysis support, M.C., E.T., A.L.P., X.F., and K.R.-H.; writing—original draft, Z.A.U. and K.R.-H.; writing—review & editing, A.U.Z., A.L.P., E.T., M.C., K.M.L., L.T.A. and K.R.-H.; funding acquisition, Z.A.U., K.R.-H.,K.M.L., and L.T.A.; resources, K.M.L., L.T.A., and K.R.-H.; supervision, K.M.L., L.T.A., and K.R.-H.

## DECLARATION OF INTERESTS

Stanford University has filed patent applications related to endothelial differentiation.

## DECLARATION OF GENERATIVE AI AND AI-ASSISTED TECHNOLOGIES

During the preparation of this work, the author used ChatGPT to assist with troubleshooting code for the analysis of bulk RNA-seq data. All content generated by the tool was reviewed and edited by the author who takes full responsibility for the content of the publication.

## SUPPLEMENTAL FIGURE LEGENDS

**Figure S1.**
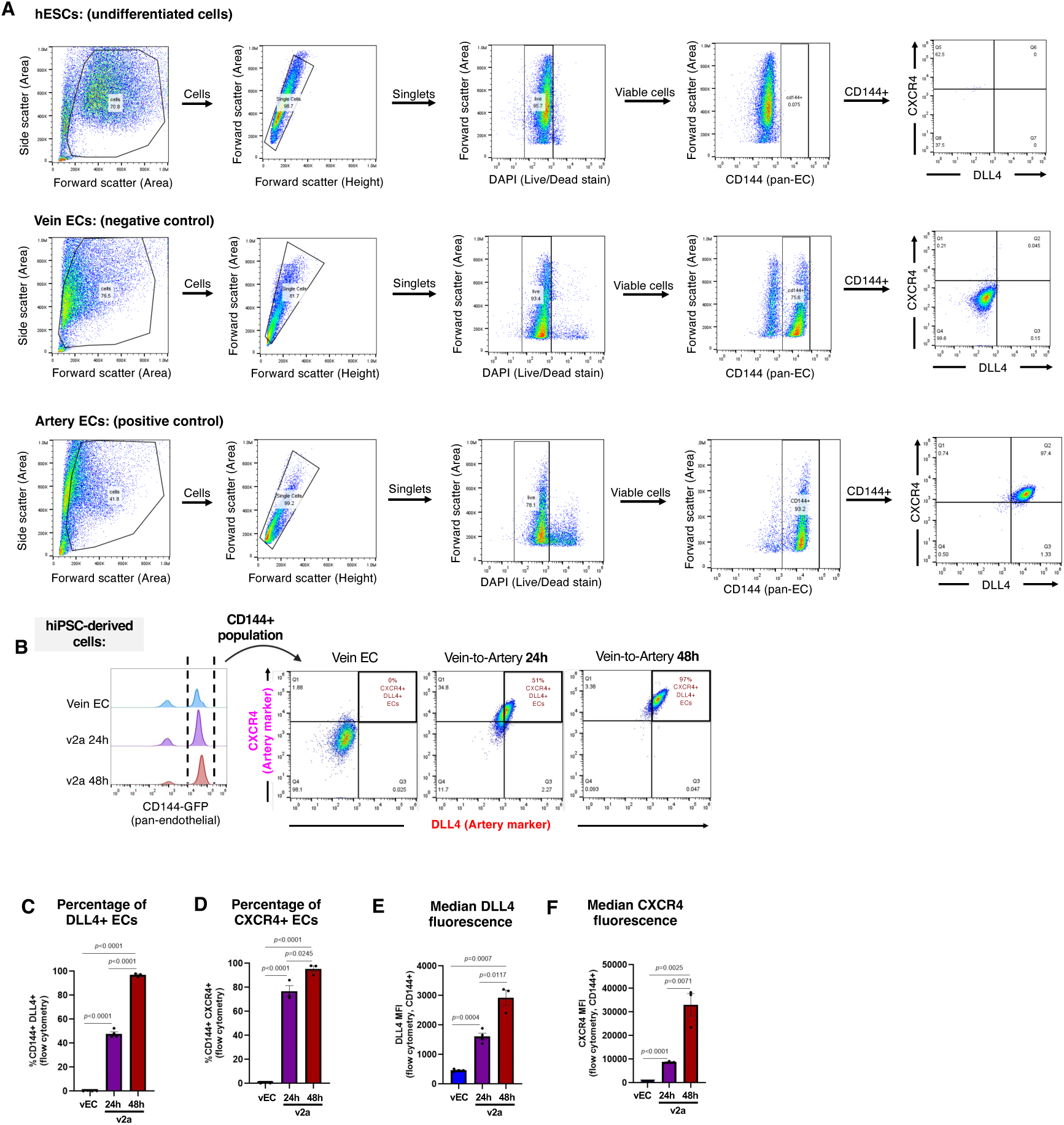
Arterial induction of hPSC-derived venous endothelial cells. **(A)** Representative gating strategy for undifferentiated human ESCs (negative for all markers). Vein ECs were used as a negative control for arterial markers (DLL4-APC, CXCR4-PE_Cy7) and as a positive control for pan-endothelial marker CD144-FITC. **(B)** Representative flow cytometry plots depicting DLL4^+^CXCR4^+^ expression within CD144^+^cells in hiPSC-derived vECs, and v2a ECs at 24 and 48 hours. **(C-F)** Quantification of the percentage of CD144^+^ ECs expressing DLL4^+^ **(C)** and CXCR4^+^ **(D)** populations in hiPSC-derived cells and their corresponding MFIs **(E, F)**, showing progressive arterial marker induction. In histograms, each dot represents one independent experiment, and error bars are mean +/- SEM. *p*-values are shown.

**Figure S2.**
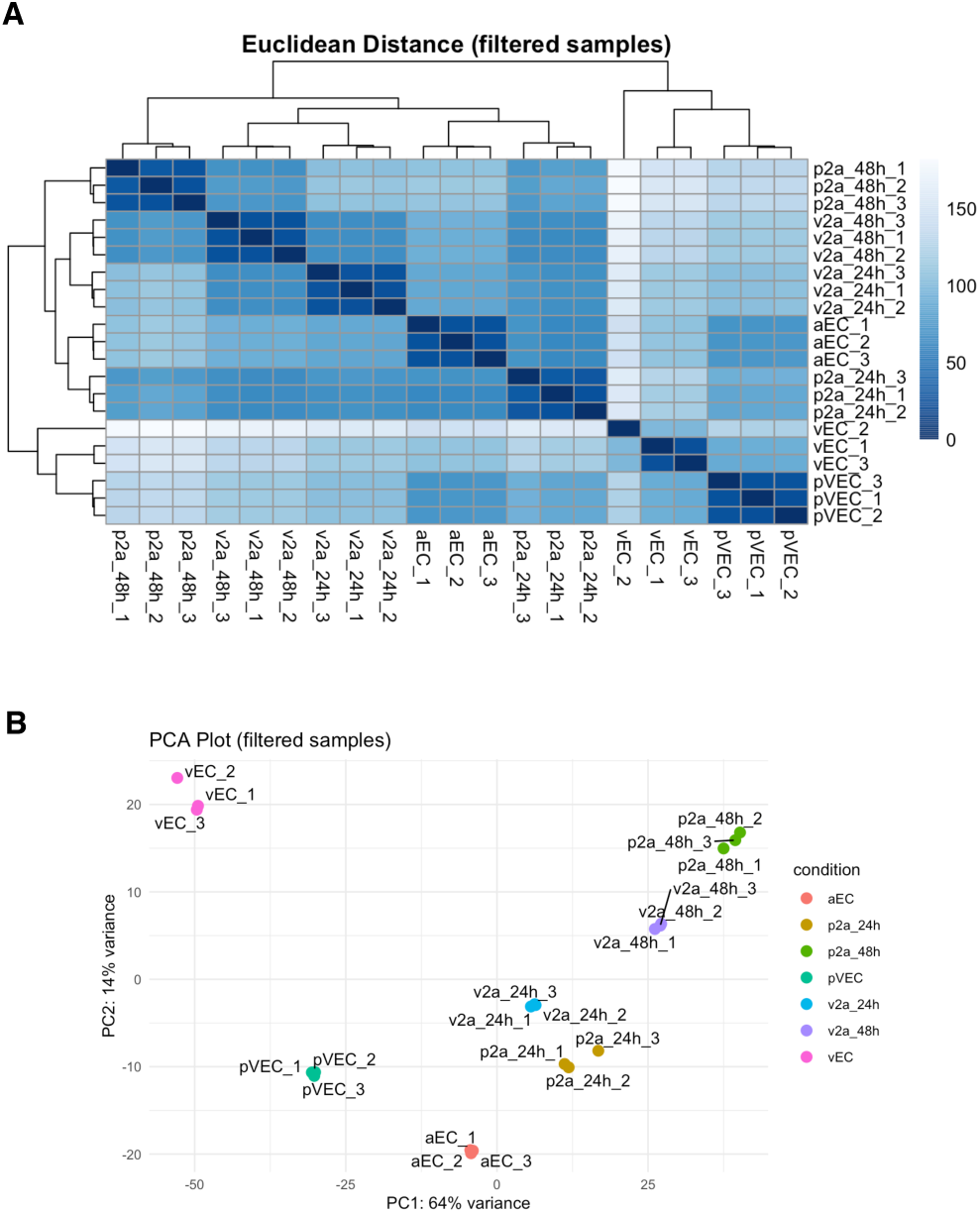
Transcriptomic relationships among hPSC-derived endothelial cells and arterialized cells. **(A)** Euclidean distance plot showing the global transcriptomic similarity across all samples, including arterial ECs (aEC), venous ECs (vEC), primed ECs (pVEC), primed EC-to-artery cells at 24 and 48 hours (p2a 24h and 48h), and vein-to-artery cells at 24 and 48 hours (v2a 24h and 48h). **(B)** Principal component analysis (PCA) plot of the indicated samples, illustrating the variance and clustering of replicates for each cell type and time point. Each dot represents one technical replicate.

**Figure S3.**
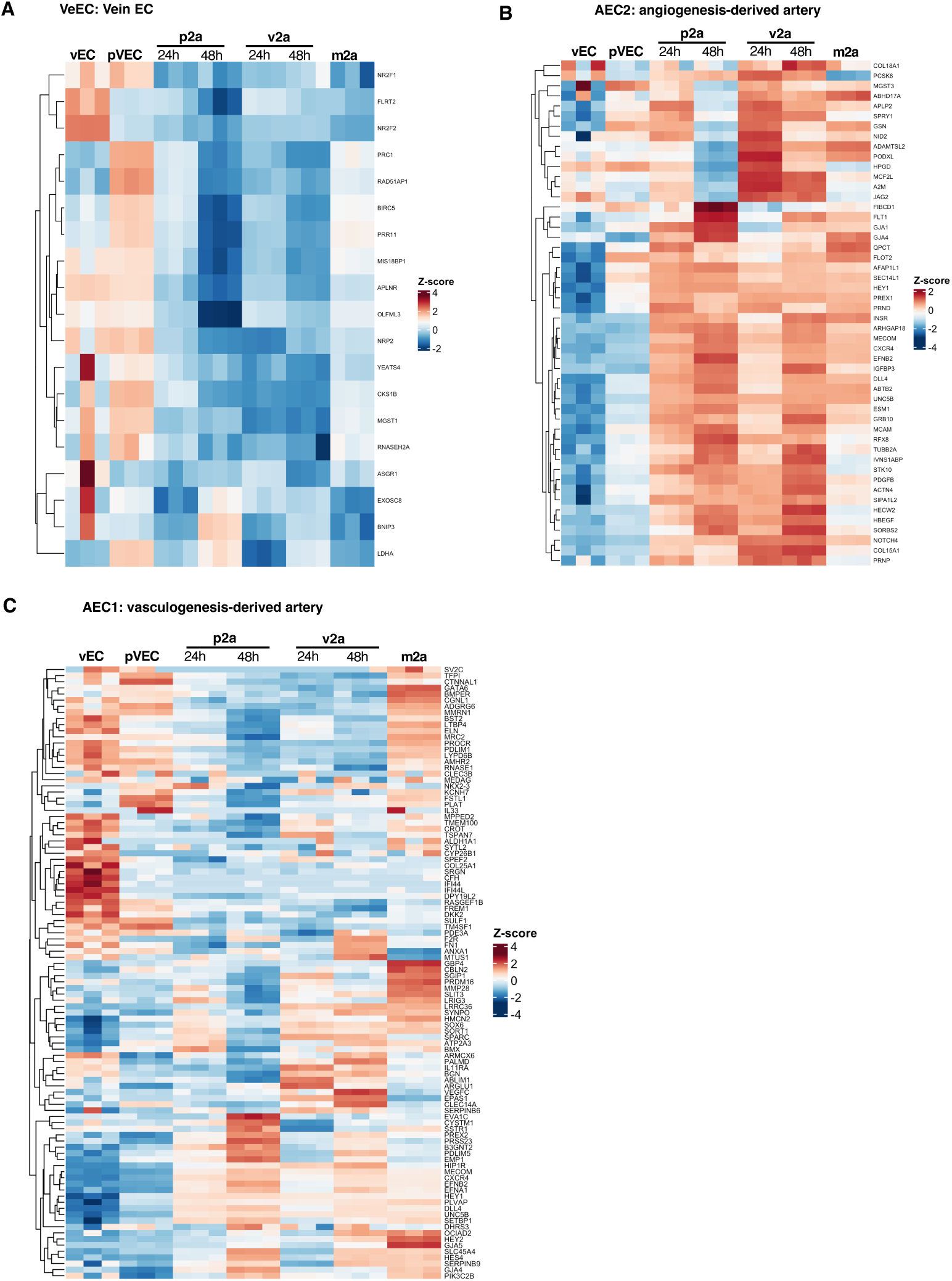
Human *in vivo* endothelial gene sets from Hou et al. applied to our *in vitro* samples. **(A-C)** Heatmaps show the expression of the top genes from three human endothelial clusters defined by Hou et al.: VeEC (vein)**(A),** AEC2 (angiogenic arteries)**(B),** and AEC1 (vasculogenic arteries)**(C),** projected onto our in vitro samples (vEC, pVEC, p2a 24h and p2a 48h, v2a 24h and v2a 48h, and m2a ECs). These heatmaps show how venous ECs arterialize and acquire angiogenic and arterial identities during the vein-to-artery endothelial transition.

**Figure S4.**
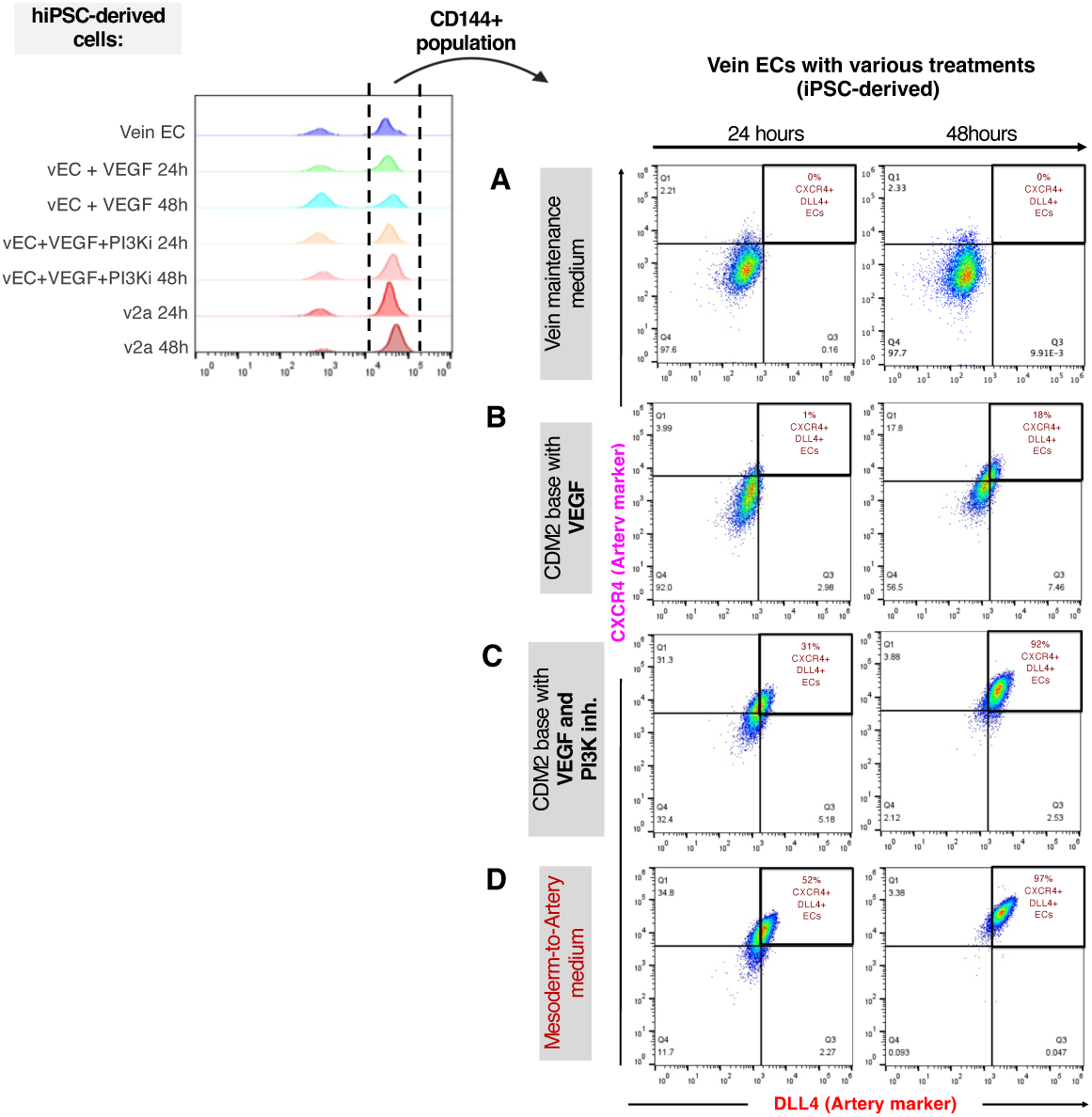
PI3K inhibition and VEGF are sufficient to arterialize vein endothelial cells in human iPSC-derived cells. **(A-D)** Representative flow cytometry plots of CD144^+^ DLL4^+^CXCR4^+^ cells across four treatment conditions.

**Table S1:**
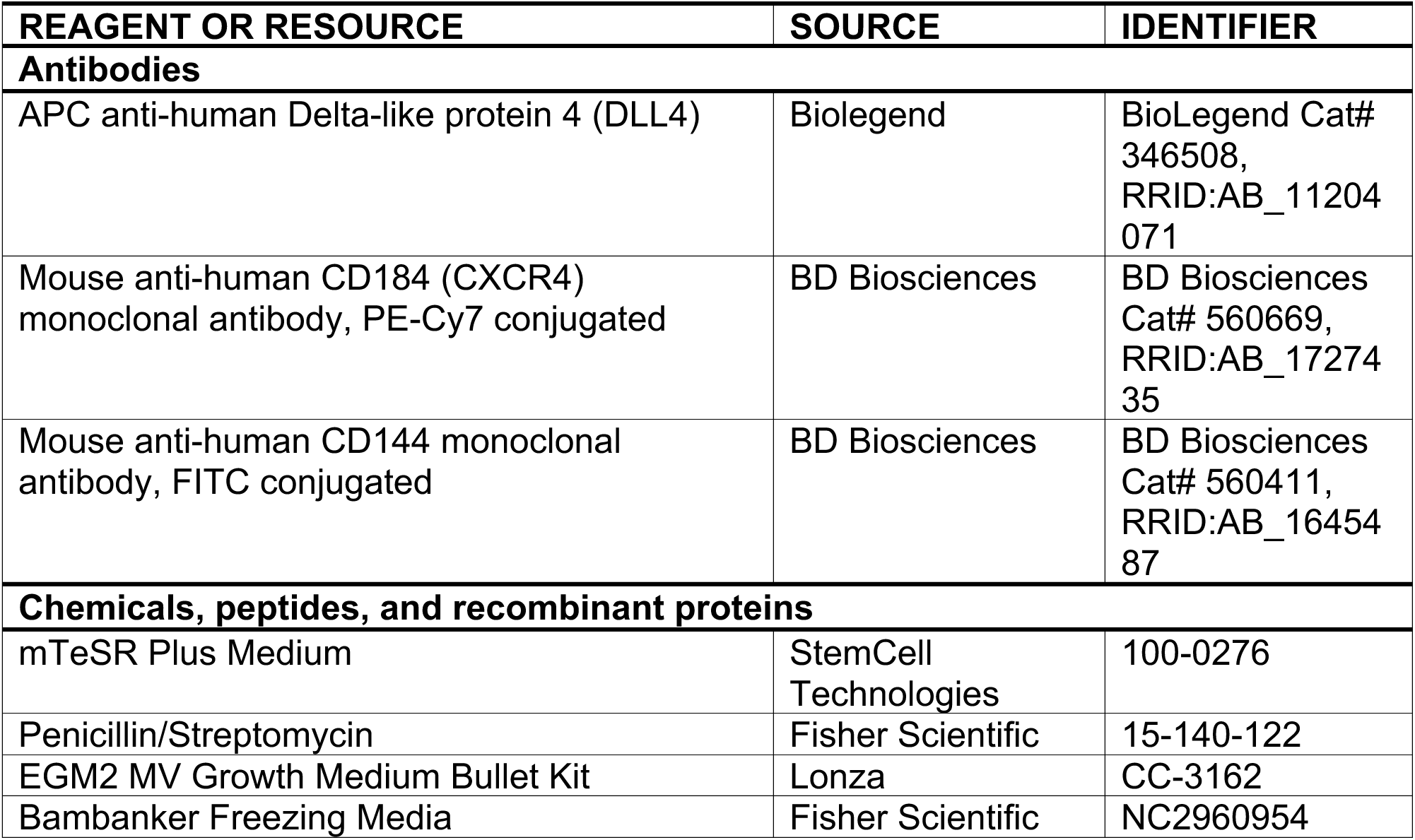
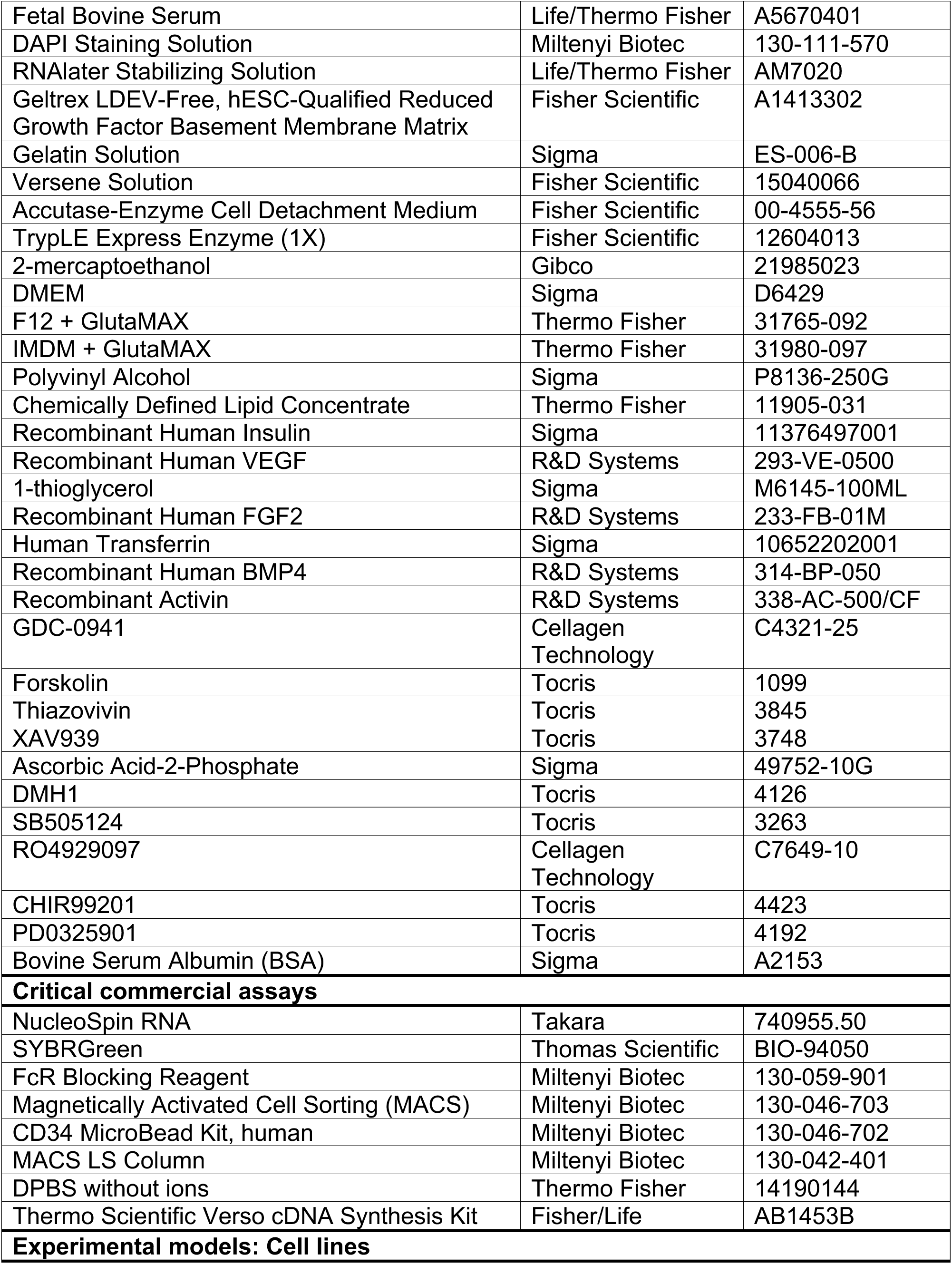
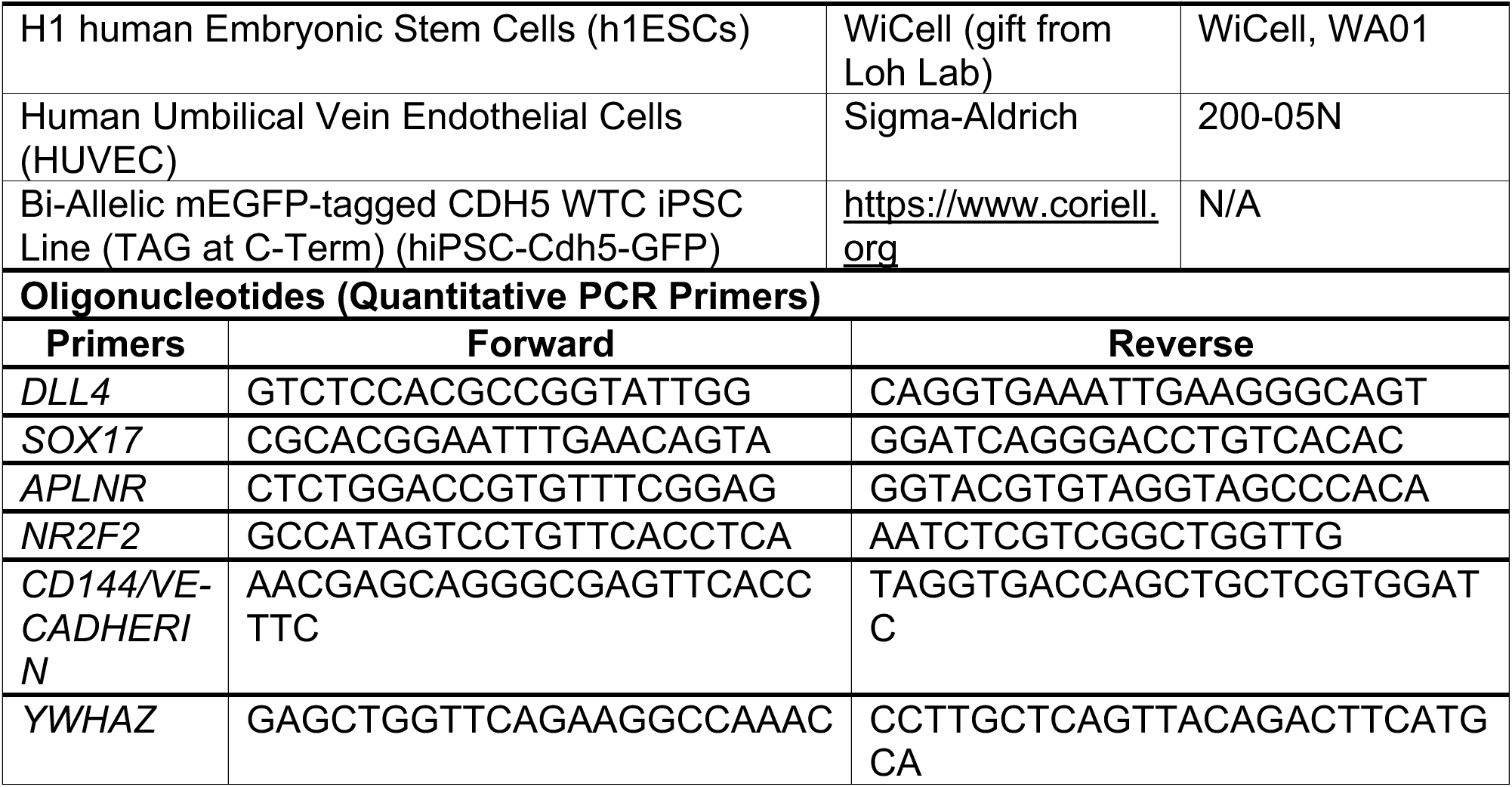
Bulk RNA-seq of hPSC-derived endothelial samples. Table of normalized raw RNA-seq counts filtered for |log2 fold change| >2.0 and adjusted p-value ≤0.05.

## MATERIALS AND METHODS

### Cell culture

All cells in this study were cultured in standard incubator conditions (20% O2, 5% CO2 and 37 °C).

### Human pluripotent stem cell lines

H1 human embryonic stem cells (H1 hESCs; WiCell), and WTC11 *CDH5-GFP* reporter human induced pluripotent stem cells (hiPSCs; Allen Institute for Cell Science/Coriell Institute) were used in this study. The biological sex of the human pluripotent stem cells (hPSCs) corresponds to that of the parental hPSC line; both lines are male.

### Human Umbilical Vein Endothelial Cells

Human umbilical vein endothelial cells (HUVEC, Lonza) were cultured on 0.1% Gelatin-coated plates in EGM2 medium (Lonza, CC-3162), with medium refreshed every 1–2 days. The biological sex of these primary human endothelial cells was not determined.

### Human pluripotent stem cell-derived endothelial cells

#### Preparing CDM2 basal medium and hPSC differentiation into artery and vein cells

Chemically defined medium (CDM2) was prepared as previously described^35^. The medium consisted of equal parts IMDM + GlutaMAX (Thermo Fisher, 31980-097) and F12 + GlutaMAX (Thermo Fisher, 31765-092), supplemented with 1 mg/mL polyvinyl alcohol (Sigma, P8136-250G), 1% v/v chemically defined lipid concentrate (Thermo Fisher, 11905-031), 450 μM 1-thioglycerol (Sigma, M6145-100ML), 0.7 μg/mL recombinant human insulin (Sigma, 11376497001), 15 μg/mL human transferrin (Sigma, 10652202001), and 1% v/v penicillin/streptomycin (Thermo Fisher, 15070-063). Polyvinyl alcohol was dissolved by gentle warming and stirring, and the medium was sterilized by filtration through a 0.22 μm filter before use.

HPSCs were differentiated into arterial (aECs), primed (pVECs), and venous (vECs) endothelial cells following established protocols (Ang *et al*., 2022, 2026). Briefly, hPSCs were first induced toward primitive streak using CDM2 supplemented with Activin A (30 ng/mL, R&D Systems), BMP4 (40 ng/mL, R&D Systems), CHIR99021(6 μM, Tocris), FGF2 (20 ng/mL, Thermo Fisher) for 24 hours. Followed by induction into dorsal lateral mesoderm (DLM) using CDM2 supplemented with BMP4(40 ng/mL), GDC-0941 (2.5 μM, Cellagen Technology), Forskolin (10 μM, Tocris), SB-505124 (2 μM, Tocris), VEGF (100 ng/mL, R&D Systems), XAV939 (1 μM, Tocris) and ascorbic acid-2-phosphate (AA2P; 200 μg/mL, Sigma) for 24 hours. To generate aECs, DLM cells were treated with CDM2 medium supplemented with CDM2 medium supplemented with Activin A (15 ng/mL), DMH1 (250 nM, Tocris), GDC-0941 (2.5 μM), VEGF (100 ng/mL), XAV939 (1 μM) and AA2P (200 μg/mL) for 24 hours. To generate primed ECs DLM cells were differentiated in CDM2 medium supplemented with SB505124 (2 μM), DMH1 (250 nM), RO4929097 (2 μM, Cellagen Technology), VEGF (100 ng/mL), XAV939 (1 μM) and AA2P (200 μg/mL) for 24 hours. Finally differentiating primed ECs in CDM2 medium supplemented with SB505124 (2 μM), RO4929097 (2 μM), PD0325901 (500 nM, Tocris), CHIR99021 (1 μM) and AA2P (200 μg/mL) for 24 hours yielded vECs.

Seeding cells at the correct density at different steps is important to subsequently achieve highly efficient differentiation into artery or vein ECs (Loh *et al*., 2025). Here, hPSCs were seeded at 190,000 – 200,000 cells per well in a 12-well plate. For aECs, the remainer of the differentiation was carried out in these wells. For vECs, the cells are re-seeded at the dorsal lateral mesoderm stage by lifting with Accutase and re-plating in primed EC media at 250,000-300,000 cells/well, as previously described (Ang *et al*., 2022). Subsequently, the primed ECs are differentiated into vein ECs, as previously described (Ang *et al*., 2022; Loh *et al*., 2025).

#### Vein or primed endothelial cell arterialization

Confluent monolayers of hPSC-derived vECs or pVECs were cultured in mesoderm-to-artery (m2a) medium for 24 or 48 hours to induce arterial specification. The m2a medium consisted of CDM2 supplemented with Activin A (15 ng/mL), DMH1 (250 nM, Tocris), GDC-0941 (2.5 μM), VEGF (100 ng/mL), XAV939 (1 μM) and AA2P (200 μg/mL)(Ang *et al*., 2022). For some experiments, one of these factors was removed from the media (Fig. 3B). For experiments in Fig. 3G, vECs were arterialized with CDM2 plus either VEGF (100 ng/mL) or VEGF (100 ng/mL) + GDC-0941 (2.5 μM). Where indicated, vECs were kept in vein maintenance media: EGM2 medium supplemented with SB505124 (2 μM) and RO4929097 (2 μM) (Ang *et al*., 2022; Loh *et al*., 2025).

#### Flow Cytometry

Undifferentiated and differentiated hPSCs were dissociated into single cells with Accutase (Fisher Scientific) (5 min, 37°C), diluted 1:5 in DMEM/F12 (Sigma), and centrifuged at 300 × *g* for 5 min at room temperature. Cell pellets were resuspended in FACS buffer (PBS [Thermo Fisher] containing 1 mM EDTA [Invitrogen] and 2% v/v FBS [Atlanta Biologicals]) and incubated with fluorescently conjugated primary antibodies—CD144-FITC (BD Biosciences), DLL4-APC (BioLegend), CD73-PE (BioLegend), and CD184 (CXCR4)-PE/Cy7 (BD Biosciences)—for 30 minutes on ice, protected from light. Cells were washed once with FACS buffer and resuspended in 500 μL FACS buffer containing DAPI (1:500; BioLegend) for live/dead cell discrimination.

Flow cytometry was performed using Sony MA900 Cell Sorter (Sony Biotechnology Inc., San Jose, CA, USA) equipped with four lasers (405, 488, 561, and 640 nm). Data were analyzed in flow cytometry analysis software, FlowJo v10.10.0, by sequential gating based on forward and side scatter, followed by height and width parameters for doublet exclusion. Live (DAPI-negative) cells were used for all marker analyses and quantification of population frequencies (see **Supplemental Fig. S1A** for gating strategy).

#### Quantitative PCR

Total RNA was extracted from undifferentiated hPSCs and hPSC-derived endothelial cells using the NucleoSpin RNA Kit (Macherey-Nagel) according to the manufacturer’s instructions. RNA was reverse transcribed into cDNA using the Verso cDNA Synthesis Kit (Thermo Fisher Scientific) following the manufacturer’s protocol. Quantitative Polymerase Chain Reaction (qPCR) was performed in a 384-well format on a QuantStudio 7 Flex Real-Time PCR System (Thermo Fisher Scientific) using gene-specific primers (Key Resources Table). Gene expression was normalized to both the housekeeping gene *YWHAZ* and the pan-endothelial marker *CD144* (VE-cadherin) to control for variation in RNA input and endothelial cell content. Relative expression was calculated using the ΔΔCT method.

#### Bright-Field Imaging of Live Endothelial Cells

Live human umbilical vein endothelial cells (HUVECs) and differentiated hPSC-derived ECs were imaged using a Leica microscope (Leica Microsystems, Wetzlar, Germany) equipped with S40/0.45 objective lens and using the Leica Application Suite (LAS) software for image acquisition. Cells were maintained in their culture medium during imaging, and images were captured using a digital camera attached to the microscope. Bright-field images were used to assess cell morphology, confluency, and overall culture health. Scale bars were added using Fiji/ImageJ. Brightfield images were used to assess cell morphology, and confluency.

## QUANTIFICATION AND STATISTICAL ANALYSIS

Statistical analyses were performed using GraphPad Prism v10. Unpaired Student’s *t*-tests were used for all comparisons, and exact *p* values are indicated.

### Bulk RNA Sequencing

Bulk RNA sequencing of hPSC-derived ECs was performed largely as described (Ang *et al*., 2022). Differentiated ECs were enriched for CD34+ cells using CD34 microbead-based Magnetically Activated Cell Sorting (MACS, Miltenyi Biotec 130-046-702) prior to RNA extraction to minimize contamination from non-endothelial populations. RNA was extracted using the NucleoSpin RNA Kit (Macherey-Nagel) and sent to Novogene for library preparation and sequencing.

Raw sequencing reads underwent quality control with FastQC (Corson *et al*., 2003) and adaptor trimming and filtering using trimmomatic (Bolger, Lohse and Usadel, 2014). Reads were aligned to the GRCh38 (hg38) reference genome (gencode v44 primary assembly annotation), and gene-level counts were quantified using HTSeq (Anders, Pyl and Huber, 2015).

Raw counts were normalized and processed using DESeq2(Love, Huber and Anders, 2014) in to identify genes differentially expressed between hPSC-derived aEC and vECs. Processing and filtering of the data frames were performed using dplyr (Wickham *et al*., 2023) and tidyverse (Wickham *et al*., 2019) and visualized using ggplot2 (*Create Elegant Data Visualisations Using the Grammar of Graphics*, no date). Heat maps, Violin plots, and Volcano plots were generated to visualize differential expression, filtering for |log2 fold change| >2.0 and adjusted p-value ≤0.05. Statistical significance for pairwise comparisons was determined using unpaired t-tests. The raw RNA-seq counts of our hPSC-derived EC samples are provided in **Table S1.**

### Artery and Vein GSVA Scoring

Artery and vein scores were generated using Gene Set Variation Analysis (GSVA) (Hänzelmann, Castelo and Guinney, 2013), implemented with GSVA Bioconductor package in R. Gene sets were derived from the *in vivo* scRNA-seq endothelial data described in Hou et al, 2022 (Hou *et al*., 2022). We extracted all genes in the vein (VeEC) and arterial (AEC1 and AEC2) endothelial clusters in human datasets. GSVA was then used to calculate sample-level enrichment scores. To visualize the data, heatmaps and bar plots were generated using the cluster-defining genes.

## RESOURCE AVAILABILITY

### Lead contact

Requests for further information and resources should be directed to and will be fulfilled by the lead contact, Kristy Red-Horse.

### Materials availability

This study did not generate new unique reagents. All materials used in this study are available from the lead contact without restriction.

### Data and code availability

All bulk RNA sequencing data generated in this study is available on NCBI Gene Expression Omnibus (GEO) under GSE312764. Normalized count matrix of all samples in our study are provided in **Table S1**. Genes were filtered according to the cutoff criteria described in the Bulk RNA-seq methods section.

## Notes

### Summary of Updates

This version of the manuscript has been revised to reflect the final accepted manuscript. Figures 1 through 4 have been updated in the main text, and Supplementary Figures S1 through S4 and their legends have been updated accordingly. Minor formatting and text clarifications were made to align with the final journal version.

